# Non-invasive tracking of hippocampal theta oscillations

**DOI:** 10.64898/2026.01.13.699218

**Authors:** E. Marcantoni, C. Daube, D. Wang, C. Cao, B. Su, S. Zhan, R.A.A. Ince, L. Parkkonen, S. Palva, D. Bush, S. Hanslmayr

**Affiliations:** Centre for Neurotechnology, School of Psychology and Neuroscience, University of Glasgow, Glasgow (UK); Department of Neuroscience, Physiology and Pharmacology, University College London, London (UK); Department of Neurosurgery, Ruijin Hospital, Shanghai Jiao Tong University School of Medicine, Shanghai (China); Department of Neuroscience and Biomedical Engineering, Aalto University, Espoo (Finland); Neuroscience Center, Helsinki Institute of Life Science, University of Helsinki, Helsinki (Finland)

## Abstract

Hippocampal theta oscillations play a crucial role in the formation of episodic memories by binding multisensory information into coherent episodes. Brain stimulation studies suggest that memory performance can be modulated by targeting these oscillations, offering a potential way of treating memory disorders. However, to effectively engage these rhythms, precise knowledge about their presence, frequency, phase and location is required, which has been a challenge, particularly for non-invasive methods such as magnetoencephalography (MEG).

Here, we first asked if hippocampal signals are detectable in MEG recordings. Using simultaneous MEG-intracranial EEG (iEEG) data, we show that invasively recorded hippocampal activity is reflected in MEG sensor signals, demonstrating the feasibility of detecting deep-brain activity non-invasively. Building on this, we introduce a MEG-based analysis pipeline to track hippocampal theta frequency over time. Applied across three independent datasets, the pipeline captured characteristic hippocampal theta frequency patterns in both rodents and humans, and MEG-derived hippocampal frequencies were consistent with those observed simultaneously in intracranial EEG.

These findings provide evidence that MEG can reliably track individual hippocampal oscillatory dynamics, paving the way for future non-invasive closed-loop interventions that adapt stimulation frequency and timing to ongoing oscillations.

## Introduction

Hippocampal theta oscillations are considered a hallmark of memory processing and have been consistently linked to the encoding and retrieval of episodic experiences ^1–4^. These oscillations organise neural firing sequences within the hippocampal circuit and provide a temporal framework that binds distributed information into coherent representations, thereby supporting the formation of new memories^5–7^.

In animal models, hippocampal theta rhythms are tightly coupled to learning and exploratory behaviour, with experimental evidence showing that the phase of theta modulates long-term potentiation (LTP) and, consequently, the strengthening of synaptic connections underlying memory formation ^8–10^.

In humans, the ability to measure and modulate hippocampal oscillations is of great interest, both for understanding the neural dynamics supporting memory and for developing interventions that target these rhythms in clinical populations. Direct evidence that rhythmic hippocampal activity supports episodic memory formation in humans comes from intracranial recordings, which have shown that hippocampal theta and gamma activity during encoding predict subsequent memory performance ^11–14^. Given this strong association, there is growing interest in understanding whether hippocampal theta can be externally modulated to influence mnemonic processing. In this context, rhythmic sensory stimulation (RSS) paradigms, which synchronise visual or auditory stimuli at certain frequencies, have emerged as promising non-invasive tools to modulate these hippocampally dependent processes. For instance, synchronising multisensory inputs at 4 Hz enhances episodic memory performance ^15–18^, presumably by entraining endogenous hippocampal theta oscillations involved in binding stimuli across modalities ^19^. Such findings suggest a causal contribution of theta rhythms to mnemonic processing and raise the possibility of non-invasive interventions to improve memory function. However, reported effects of rhythmic stimulation have been variable across studies ^20,21^, highlighting the need for a better characterisation of hippocampal oscillations before designing effective interventions. For instance, human hippocampal theta typically occurs as transient bouts that vary across encoding and retrieval periods ^11,22,23^. Accurately characterising when and how these oscillatory bouts occur is therefore critical for establishing reliable markers of hippocampal engagement and for guiding stimulation strategies that adapt to the brain’s ongoing state and target the oscillation when present.

One of the major obstacles to developing such methods is that recording hippocampal oscillations non-invasively remains technically challenging. Magnetic or electrical signals from deep brain structures are strongly attenuated by the distance from the sensors or spatially smeared by overlying tissue, making their reconstruction from surface neurophysiology inherently difficult. In addition, cortical activity couples more strongly to the sensors and can therefore dominate the measured signal, effectively masking contributions from deeper sources^24^. Although the feasibility of detecting hippocampal activity with magnetoencephalography (MEG) is a subject of continuous debate, the hippocampus is a particularly promising candidate. Despite its depth, the geometrical configuration of hippocampal pyramidal neurons leads to larger local field potentials than the neocortex, which may plausibly contribute to extracranial magnetic fields ^25–27^. Consistent with this view, growing evidence from simultaneous invasive and non-invasive recordings (the ground truth for deep-source detection) indicates that hippocampal signals can, under suitable conditions, be detected and reconstructed using advanced source-modelling approaches ^28–32^. Yet, validation remains limited because such simultaneous recordings are confined to patient populations and are rare.

In the present work, we provide support to the body of evidence suggesting that hippocampal oscillations can be detected non-invasively with MEG, and we develop and validate a MEG-based analysis pipeline capable of tracking hippocampal theta frequency over time.

## Results

### Assessment of the hippocampal contributions to MEG

As the first step, we asked whether hippocampal activity can be detected in MEG sensor data. To address this question, we analysed simultaneous MEG and intracranial EEG recordings from two epilepsy patients at rest (six runs in total). We applied canonical correlation analysis (CCA)^33^ in a data-driven manner to identify linear combinations of MEG sensors whose activity covaried with hippocampal iEEG signals over time, thereby testing for shared neural activity between the two modalities without making prior anatomical assumptions.

To assess the anatomical origin of these sensor-level components, we next localised the MEG projections of the CCA components in source space. Specifically, we used an LCMV beamformer to reconstruct voxel-wise MEG time series and quantified, for each voxel, the correspondence between its activity and the CCA–derived MEG component time course. This procedure revealed spatial maps with a sharp hippocampal focus (Fig. 1a), although performance varied across participants and runs. To quantify spatial specificity, we examined the correspondence between beamformed time series from all voxels and the CCA projections as a function of distance to the hippocampus for each run. Ideally, this correspondence would be expected to peak at hippocampal locations and gradually decline with increasing distance. In line with this, we observed negative correlations between correspondence strength and distance from the hippocampus, indicating that the strength of MEG–iEEG correspondence decreased with increasing distance from hippocampal regions (Fig. 1b).

**Fig. 1.**
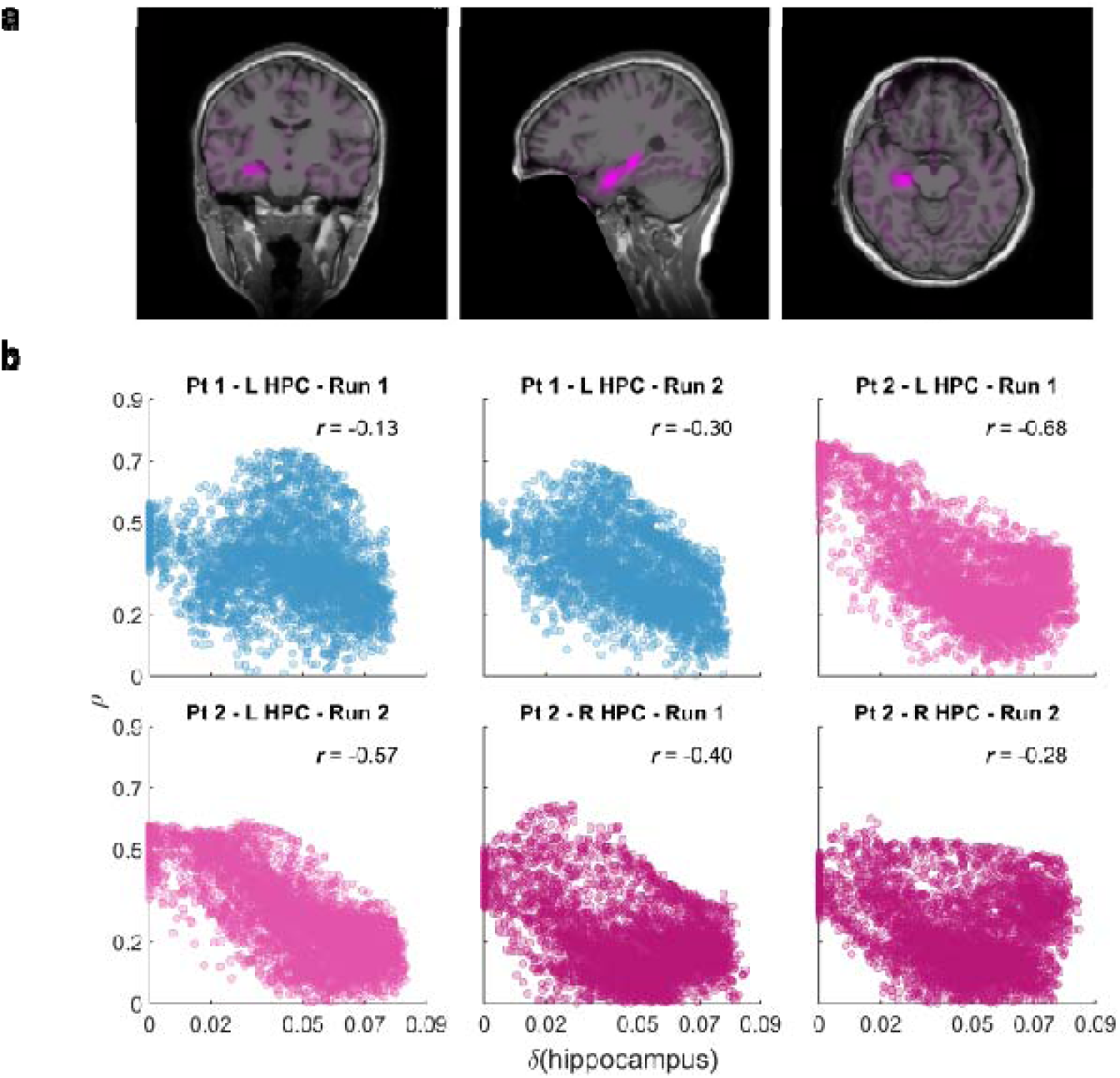
Assessment of the hippocampal contributions to MEG. **a** Example from one participant (Patient 2 - left hippocampus; best performing run) illustrating the source localisation of data-driven CCA filters. Canonical Correlation Analysis is applied to identify MEG components that are maximally correlated over time with hippocampal iEEG, with no a priori constraints about frequency, timing, or anatomical location. The cross-validated prediction performance, projected into source space, peaks in the hippocampus (magenta overlay). **b** CCA performance (best component) as a function of distance from the hippocampus, shown separately for each participant and run. Each scatterplot displays all source-space grid points; the x-axis reflects the Euclidean distance to the nearest hippocampal voxel (δ) in metres, and the y-axis shows the cross-validated model performance (*p*). Numbers report the Pearson correlation between δ and *p*, used as a specificity index (negative values indicate higher performance closer to the hippocampus). For each run, results for the best CCA component are shown.

However, a much more ubiquitous situation is that researchers only have access to MEG recordings, from which they will estimate activity from hippocampal sources using inverse models. We therefore assessed how well hippocampal signals could be recovered in this typical scenario and compared the performance to the CCA approach. Specifically, we applied a cross-correlation analysis to quantify the temporal correspondence between hippocampal iEEG signals and MEG hippocampal source estimates reconstructed using an LCMV beamformer (see Methods). The resulting spatial maps were more diffuse than those obtained with CCA but still showed elevated correspondence near the hippocampus (Supplementary Fig. 2a). Despite this reduced focality, the cross-correlation approach nonetheless preserved the expected relationship between performance and anatomical distance: across runs, model performance decreased as the distance from the hippocampal region increased (Supplementary Fig. 2b).

To directly compare the two approaches, we summarised each run using two metrics: the spatial specificity metric, expressed as the sign-reversed correlation between voxel-wise MEG–iEEG correspondence and anatomical distance to the hippocampus (such that higher values indicate greater hippocampal specificity), and a measure of sensitivity. Sensitivity quantifies the overall strength of the MEG-iEEG correspondence and was defined as the mean cross-validated correlation at the best-performing solution (the best grid point of the best electrode for cross-correlation, or the best component for CCA). This comparison revealed similar performance across methods (Supplementary Fig. 2c), with only a slight trend favouring CCA. Across runs, mean specificity was 0.36 ± 0.29 (mean ± SD) for cross-correlation and 0.39 ± 0.20 for CCA, while mean sensitivity was 0.22 ± 0.10 and 0.28 ± 0.16, respectively.

Taken together, these analyses demonstrate that hippocampal theta activity recorded with intracranial electrodes is detectable by MEG sensors: in both approaches, correlations were highest near hippocampal locations and declined with distance. At the same time, the direct comparison between methods shows that while a data-driven approach that constructs the MEG spatial filters using iEEG data (CCA) works slightly better, a classic source localisation approach that estimates hippocampal activity without access to iEEG data achieves comparable performance.

### Pipeline validation

Having demonstrated that hippocampal theta oscillations can be captured with MEG, we then developed an MEG-based analysis pipeline to track hippocampal theta dynamics over time. Specifically, we focused on detecting when individual theta bouts occur and estimating their frequency (Fig. 2a; see Methods for full details). First, we reconstruct hippocampal activity with a spatial filter from the MEG sensor signal using a Linearly Constrained Minimum Variance beamformer spatial filter (LCMV)^34^. Next, we enhance the signal-to-noise ratio of theta activity by deriving components that emphasise rhythmic fluctuations in the theta band relative to broadband activity using generalized eigendecomposition (GED)^35^. Finally, we separate genuine from spurious oscillatory bursts in the resulting time course and estimate their centre frequencies using the cyclic homogeneous oscillation method (CHO)^36^.

**Fig. 2.**
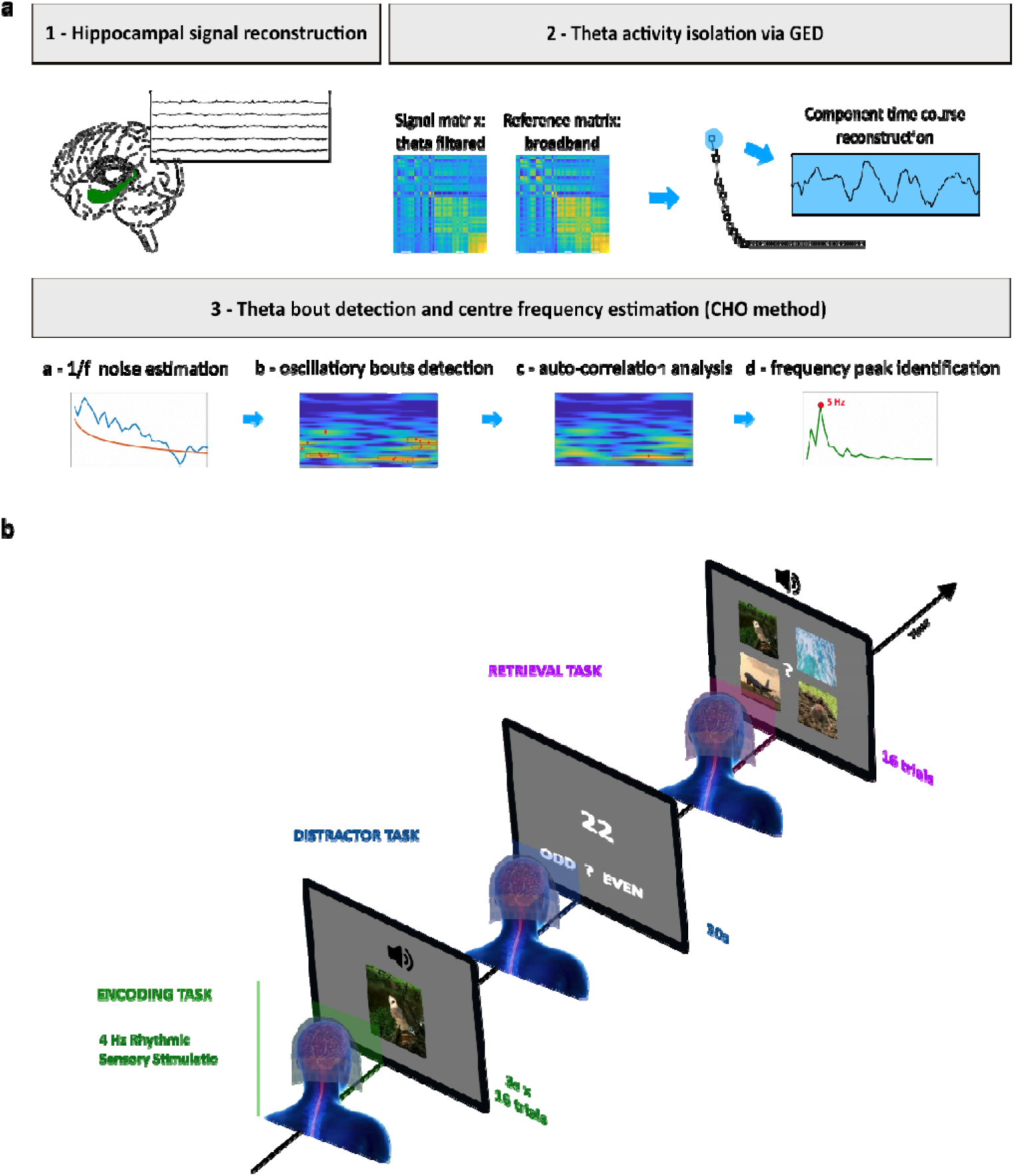
MEG processing pipeline and task structure of validation dataset 2. **a** MEG pipeline: (1) Hippocampal signals are extracted via source modeling (LCMV beamforming); (2) generalized eigenvalue decomposition (GED) is used to enhance theta-range oscillatory components; (3) theta bouts are identified and their centre frequencies estimated with the CHO method through (a) 1/f separation, (b) bouts detection, (c) autocorrelation, and (d) peak-frequency estimation. **b** Task design on the validation dataset 2: 4 Hz RSS MEG paradigm. During encoding (16 trials per block), unrelated video-sound pairs flickering and fluttering at 4Hz were presented for 3s, after which participants rated how well the sound suited the video. The pair could be presented either in-phase (0° phase offset) or out-of-phase (180° phase offset). During the 30s distractor task, participants judged whether numbers were odd or even. During retrieval (16 trials), each previously heard sound (3 s) was presented with four still-frame video options, and participants were asked to select the paired video (blocks n = 12).

To validate its performance, we applied the pipeline to three independent electrophysiology datasets: invasive rodent recordings during navigation (*N* = 3 rodents; 64 recordings), human MEG recordings during rhythmic sensory stimulation (*N* = 24 healthy participants), and simultaneous MEG-iEEG recordings from epilepsy patients (*N* = 2; 6 recordings).

Applying the pipeline to a rodent dataset before applying it to MEG data allowed us to verify whether the procedure implemented for detecting genuine theta oscillatory activity replicated well-established hippocampal theta dynamics, i.e. an increase in theta frequency with running speed. To this end, we applied the frequency identification steps of the pipeline to invasive rodent recordings during locomotion and correlated it with running speed. We then applied the complete pipeline to a human MEG dataset recorded during an associative memory task and during 4-Hz rhythmic sensory stimulation, assessing its ability to track hippocampal theta frequency and its presumed entrainment to an external rhythm. Finally, we applied the pipeline to the simultaneous MEG-iEEG dataset to compare the frequencies identified in the source-reconstructed signal with the ground-truth frequencies derived from iEEG recordings.

### Rodent dataset

In the first dataset, we applied part of the pipeline to intracranial recordings from rodents navigating in an open field (*N* = 3 rodents; 64 recordings). The scope of this analysis was to verify that our approach for enhancing theta activity (GED) and identifying genuine oscillatory frequencies (CHO) could be trusted. In rodents, hippocampal theta frequency is tightly linked to locomotion speed^37–39^, providing a well-established benchmark for such validation.

We observed a positive association between theta frequency and running speed across 1-s analysis windows (Fig. 3a; Pearson’s *r* = 0.27). To formally assess the relationship while accounting for repeated measurements within animals and sessions, we fitted a linear mixed-effects model with random intercepts for animal and session. This model confirmed a robust positive effect of running speed on theta frequency (β = 3.49 cm^-1^, SE = 0.059).

**Fig 3.**
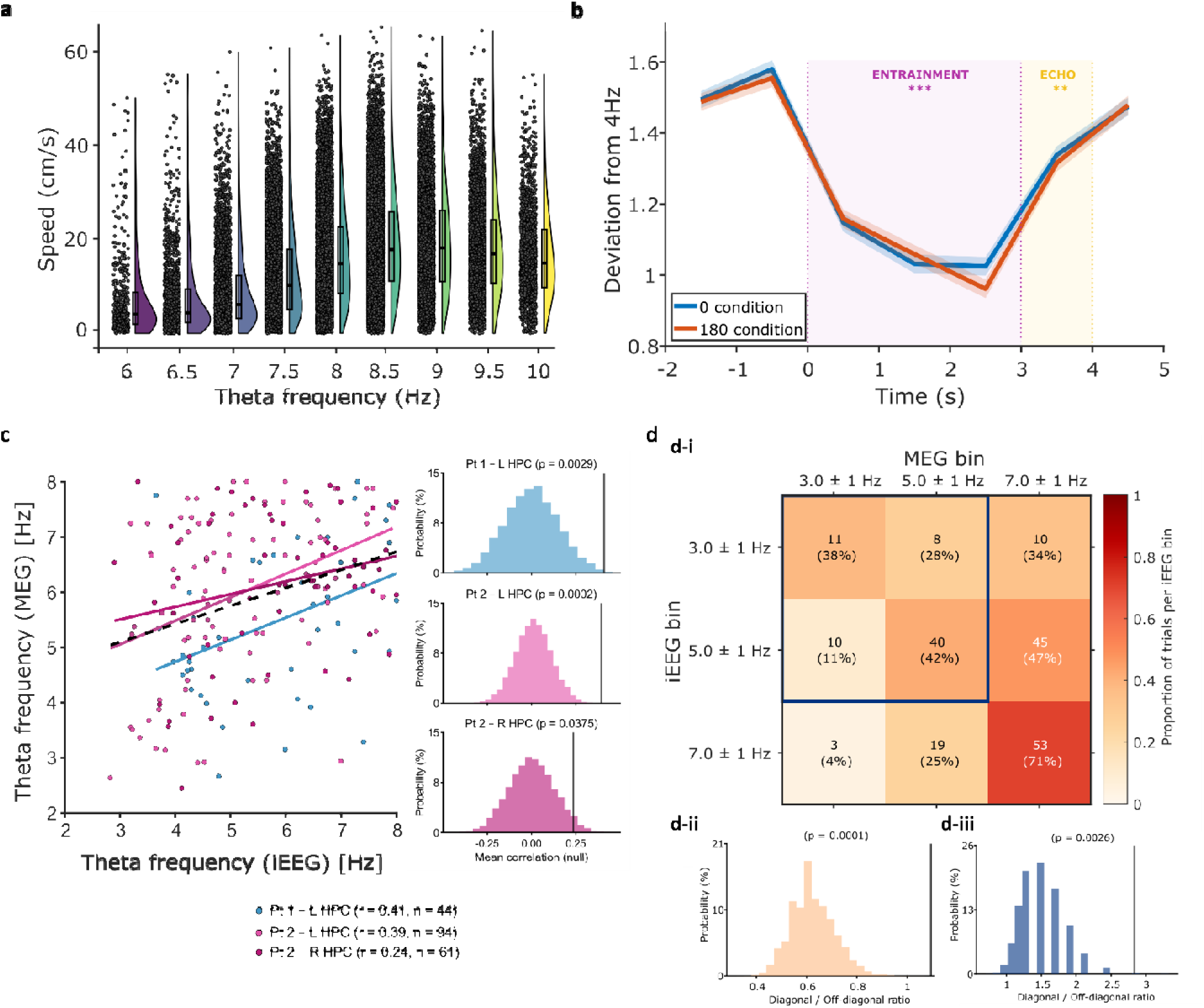
MEG pipeline validation results. **a** Distribution of rodent speed across theta frequency values. Violin plots show the density of single-trial speed observations (cm/s) at each theta frequency (Hz); black dots represent individual epochs, and boxplots indicate medians and interquartile ranges. **b** Deviation of the identified frequency from 4 Hz (identified - stimulation frequency) across seven 1s bins for 0° (blue) and 180° (red) phase-offset trials (time 0 = pair onset). Lines show means; shaded areas show standard deviation. Significant differences between stimulation and pre/post windows are marked in violet (entrainment), and those between pre-stimulus and the first post-stimulus second in yellow (entrainment echo). **c** *Left*: Scatterplot showing the correlation between theta frequencies estimated from source-reconstructed MEG data (y-axis) and intracranial EEG (iEEG) recordings (x-axis). Each data point represents a theta frequency estimate from a single epoch, with values derived independently from both iEEG and source-reconstructed MEG data. Each coloured line represents a within-hemisphere fit. The dashed black line indicates the group-level linear regression fit (shaded bands = 95% CI). *Right:* Patient-hemisphere specific observed correlations compared to null distributions (n = 10000 permutations) of correlated coefficients obtained by randomly permuting the MEG frequency vector. **d** *d-i*: Confusion matrix showing how well MEG-derived theta frequencies matched the iEEG theta frequencies (2-8 Hz). Rows show the iEEG bins and columns show the MEG bins. Values are row-normalised (proportion of trials per iEEG bin), with count and percentage overlaid in each cell. Strong diagonal structure indicates that MEG theta estimates preserve the underlying iEEG frequency structure. Dark-blue bounding box indicates the subset of bins included in the control analysis (d-iii). *d-ii:* Significance was assessed via run-wise permutation of MEG labels using a diagonal/off-diagonal ratio statistic (10,000 iterations). *d-iii:* Same analysis restricted to the lower-frequency quadrant (3 ± 1 Hz and 5 ± 1 Hz bins), excluding the highest-frequency bin.

### 4-Hz RSS MEG dataset

In the second dataset, we tested the pipeline on human MEG recordings collected in our lab (*N* = 24 healthy participants) during an associative memory task with 4-Hz flickering audiovisual stimulus pairs to be encoded and later remembered (Fig. 2b). Previous studies have shown that such rhythmic sensory stimulation can entrain memory-related theta activity in the hippocampus and improve performance when the audio and the video are presented such that they synchronize visual and auditory cortices, compared to out-of-synchrony conditions ^15,16^. This paradigm thus offered the opportunity to ask whether our method could detect RSS effects at the hippocampal level. Specifically, we hypothesised that hippocampal frequency estimates would align to 4 Hz during the rhythmic window compared with pre- and post-stimulation periods.

We found that hippocampal frequency estimates shifted toward the stimulation frequency during rhythmic sensory stimulation (Fig. 3b). A repeated-measures ANOVA revealed that the deviation from 4 Hz (Δf = identified frequency – 4 Hz) varied significantly across trial time (main effect of Time: *F*(6,138) = 64.58, *p* < 0.001, ges = 0.54). Planned contrasts showed that during stimulation (time bins 3–5), the estimated frequencies were significantly closer to 4 Hz than in the pre- (time bins 1–2; *t*(23) = −17.82, *p* < .001) and post-stimulation periods (*ps* < .001; see Supplementary Table 1). Notably, the effect persisted briefly after stimulation, with the first post-stimulation time bin (1 s after stimulus offset) still differing from the pre-stimulation baseline (time bin 6; *t*(23) = −5.94, *p* < .001), before returning toward baseline by the final post-stimulation bin (time bin 7; *t*(23) = −1.79, *p* = .074; FDR-corrected). No main effect of phase (0° vs 180°) and no Condition × Time interaction were observed (*ps* > .9).

To determine whether the observed 4-Hz alignment was hippocampus specific, we tested the alternative hypothesis that it reflected widespread responses to the RSS dominating the sensor-level signal ^40^, that could leak into deep-source estimates ^41^. We therefore examined two regions expected to entrain less to this stimulation: the primary motor cortex (M1) and dorsolateral prefrontal cortex (DLPFC). No comparable frequency shift was observed in these areas (see Supplementary Methods and Results for full analysis; Supplementary Fig. 1).

### MEG–iEEG dataset

In the third dataset, we analysed simultaneous MEG–iEEG recordings from epilepsy patients at rest (*N* = 2; 6 recordings). This dataset provided a unique opportunity to directly compare hippocampal activity reconstructed from MEG with invasive recordings from electrodes implanted in the hippocampus. Here, we asked whether the frequencies estimated from MEG would correlate with those measured directly from iEEG, providing a critical validation of the pipeline against a ground-truth reference.

Runs from the same hemisphere were combined to obtain a single MEG–iEEG correlation per hemisphere, as they were recorded from the same implanted hippocampal contacts. For each subject-hemisphere dataset, MEG- and iEEG-derived theta frequencies showed consistent positive relationships (Fig. 3c, *left panel*). Statistical significance was evaluated using a run-wise permutation test, in which MEG-derived frequency values were shuffled independently within each run to preserve run-specific characteristics before being combined again at the hemisphere level. All datasets showed significant MEG–iEEG correlations (*p* < 0.05; Fig. 3c, *right panel*).

To further assess the correspondence between MEG and iEEG, we constructed a confusion matrix by binning frequency estimates into 2-Hz intervals spanning 2–8 Hz. Each cell in the matrix reflects the proportion of trials for which a frequency in each iEEG bin was matched by a frequency in the corresponding MEG bin. The resulting distribution showed the highest values along the diagonal, indicating that frequencies estimated from MEG tended to fall within the same theta bin as those measured invasively (Fig. 3d-i).

To quantify this alignment, we computed the ratio of diagonal to off-diagonal trial counts and evaluated its significance using within-run permutations of the MEG frequency labels. The observed ratio exceeded the 99^th^ percentile of the null distribution (*p* < 0.01), confirming that the alignment between MEG and iEEG frequency estimates was unlikely to arise by chance (Fig. 3d-ii).

Because the highest frequency bin contained more trials than the others, we repeated the analysis after excluding this bin (i.e., restricting the matrix to 2-6 Hz). The diagonal/off-diagonal ratio remained highly significant (*p* < 0.01), indicating that the effect was not driven solely by the upper frequency range (Fig. 3d-iii).

## Discussion

The present study tries to take a step towards solving a methodological challenge in human neuroscience: a better and more precise characterisation of hippocampal theta frequency, a prerequisite for understanding and, ultimately, manipulating memory-related rhythms in humans. Much of what is known about hippocampal theta in humans comes from iEEG recordings in clinical populations. Although iEEG offers direct access to hippocampal activity, clinically driven electrode placement and potential confounds from pathology or medication limit generalizability ^42,43^. As a result, our current understanding of hippocampal theta in humans, its frequency variability, temporal dynamics, and engagement during cognitive states, remains fragmentary and limited. Advancing our understanding of hippocampal dynamics in healthy populations and how they can be influenced with non-invasive stimulation requires methods that can reliably detect hippocampal activity from surface recordings. Such recordings, however, face two major obstacles: limited sensitivity to deep sources and the simplifying assumptions of conventional spectral analyses. Standard methods model neural signals as stationary sums of sinusoids and typically neglect the aperiodic background. This simplification can blur the distinction between genuine rhythmic activity and spurious spectral peaks ^44^, potentially mischaracterising non-sinusoidal waveforms such as the sawtooth-shaped hippocampal theta commonly observed in invasive recordings ^45,46^. Furthermore, these methods often summarise spectral characteristics across long epochs, providing limited temporal precision.

Our work addresses these issues by combining MEG beamforming with a waveform-agnostic, time-resolved characterisation of theta occurrence. The resulting analysis pipeline first localises hippocampal activity from MEG using a spatial filter informed by anatomical priors and then detects transient theta bouts directly in the reconstructed time series. In doing so, it avoids assumptions of sinusoidal signals and provides information about when oscillatory bouts emerge, dissipate, or change in frequency.

Using this approach, we demonstrate that hippocampal signals can be detected and characterised non-invasively with MEG. Across three datasets, we were able to estimate hippocampal theta frequency with sufficient precision to reconstruct known and physiologically meaningful patterns. For instance, in rodents, theta frequency scaled with running speed, reproducing a canonical behavioural signature.

In the human 4-Hz RSS paradigm, we observed a systematic shift of hippocampal frequency estimates toward the stimulation frequency. Our findings are consistent with previous demonstrations of hippocampal involvement in this task ^16,19,47^. Indeed, the hippocampus is an area where all sensory streams are integrated to form a coherent memory representation to be encoded. This integration relies on plasticity mechanisms that are influenced by the timing of ongoing theta oscillations. Rhythmic sensory input is thought to entrain cortical areas and shape the temporal structure of the information they transmit to the hippocampus. When inputs arrive at the hippocampus at similar phases on successive theta cycles, the probability of inducing synaptic plasticity and binding these elements together is increased. This idea is consistent with invasive human work showing that theta-phase organisation of neuronal firing in the hippocampus supports successful encoding through phase-dependent plasticity mechanisms^13^ and also with recent MEG work showing that theta activity in the hippocampus is modulated by the temporal structure of audiovisual input and reinstated during memory replay^47^.

A central question is whether the observed shift reflects genuine entrainment or merely periodic evoked responses, a distinction that remains debated^48,49^. One key observation in favour of entrainment here is the persistence of an entrainment echo after stimulation offset, which is difficult to reconcile with a purely evoked-response account. One potential concern is that the observed frequency shift and echo might reflect a methodological artefact, such as temporal smearing induced by filtering. However, this is unlikely: the analysis was performed in 1-second windows treated independently, so any filter smearing would be confined within each window and thus could not create systematic differences between pre-, during-, and post-stimulation periods. Another plausible concern is leakage from widespread sensory responses, as rhythmic stimulation elicits strong entrained activity that can leak into deep-source estimates. However, we did not observe a comparable frequency shift in control regions (M1, DLPFC), arguing against leakage as the primary explanation.

Finally, in simultaneous MEG–iEEG recordings, theta frequencies estimated from MEG correlated with those measured directly from hippocampal contacts, providing ground-truth confirmation that the reconstructed signals reflect genuine hippocampal activity rather than noise or superficial neocortical leakage.

Several methodological constraints should nevertheless be acknowledged. The MEG–iEEG validation dataset consisted of only six recordings from two patients and was collected during rest, a condition in which hippocampal theta is sparse and less coherent. This likely contributed to the reduced number of detectable theta bouts and the modest cross-modal correlations. Moreover, electrode coverage within the hippocampus varied across sessions, and the intracranial recordings themselves sampled only limited portions of the structure. Consequently, even the iEEG “ground truth” only provides a spatially restricted view of hippocampal activity. Another limitation is that in our implementation, the LCMV beamformer used a “max-power” parameter, selecting the dipole direction that maximised output power. This optimisation enhances sensitivity but may not correspond exactly to the physical orientation of the underlying hippocampal generators. Because the spectral content of the reconstructed signal depends on this orientation, small discrepancies between the theta frequency estimated from MEG and that recorded intracranially are expected. These differences likely reflect geometric and spatial-filtering effects rather than physiological divergence between modalities. Furthermore, while canonical correlation analysis (CCA) identified shared MEG–iEEG components with clear hippocampal localisation in several cases, performance varied across runs, likely reflecting differences in signal quality, electrode placement, or individual variability in hippocampal coupling. Despite these limitations, it is noteworthy that a reliable correspondence between MEG and iEEG signals could still be retrieved, considering that the two modalities reflect the same underlying synaptic activity but capture it through different physical measurements, which differ substantially in their signal-to-noise ratios and spatial sensitivities. Finally, the parameter space of the proposed pipeline is necessarily broad. Choices regarding beamformer, detection thresholds, and window lengths were made to balance sensitivity and robustness under realistic constraints, but further optimisation could improve accuracy and generalisability.

Together, our results establish that we can reconstruct some features of hippocampal theta oscillations, such as frequency, non-invasively, with reasonable spatial and temporal accuracy. Although the present implementation focused only on hippocampal theta frequency, the pipeline is generalisable to other oscillatory features and processes in other deep-brain structures. Importantly, even if the current approach is not yet optimal, it may offer a better framework for investigating deep-brain oscillatory dynamics and for guiding stimulation protocols beyond conventional fixed-frequency methods. This approach thus opens new avenues for studying memory-related oscillations and for developing closed-loop stimulation paradigms that align external inputs with the brain’s ongoing rhythmic states. Such precision characterisation of hippocampal theta could ultimately inform non-invasive interventions to restore or enhance memory function in clinical populations.

## Methods

### MEG Pipeline

We developed a pipeline to be able to detect the presence of theta oscillations in the hippocampus from MEG signals. The pipeline consisted of three main steps: hippocampal source reconstruction, enhancement of theta-band activity using generalized eigendecomposition (GED), and identification of transient theta oscillations using the cyclic homogeneous oscillation (CHO) method, whose parameters were adapted to the specific dataset analysed (Fig. 2a):

1. **Hippocampal source reconstruction**. Source activity was reconstructed using a linearly constrained minimum variance (LCMV) beamformer ^34^. Subject-specific forward models were constructed from T1-weighted MRI. Cortical and subcortical structures were segmented using FreeSurfer^50^ (v7.4.1; https://surfer.nmr.mgh.harvard.edu/). These outputs were imported into MNE-Python for MEG forward modelling. A single-shell boundary element model (BEM) based on the FreeSurfer-derived inner skull surface was constructed, and MEG–MRI coregistration was obtained using digitised fiducials and scalp points; the resulting transform aligned the head coordinate frame to MRI. The source space combined: (i) a sparse cortical surface (e.g., oct2–oct3 spacing) and (ii) volumetric hippocampal sources defined from the subject-specific segmentation (*aseg.mgz*) generated by FreeSurfer (Left/Right-Hippocampus). The lead fields were then computed for the combined source space, with a minimum inner-skull distance of 5 mm. This process creates a gain matrix mapping each dipole to the MEG sensors. LCMV spatial filters were computed from the subject-specific forward model together with noise and data covariance matrices. The data covariance matrix was computed from the experimental data over the entire trial time window and used to construct a single set of spatial filters. The noise covariance matrix, used for sensor whitening, was estimated from empty-room recordings processed identically to the experimental data for the 4-Hz RSS MEG dataset. For the MEG–iEEG dataset, in which empty-room recordings were unavailable, noise covariance was estimated using an ad hoc empirical estimator provided in MNE-Python (using the function “*mne.make_ad_hoc_cov*”). To avoid noise amplification when combining magnetometers and gradiometers, the effective sensor rank was derived from the whitening covariance and passed to the inverse solution^51^. LCMV filters were constructed using a regularisation parameter λ ≈ 0.05 (fraction of average sensor power), unit-noise-gain (weight normalisation), and orientation selection set to max-power per location. The resulting spatial filters were applied to the data to obtain source-level time series across the mixed cortical-hippocampal source space, after which only the hippocampal sources were retained for subsequent analyses.
2. ***Theta activity enhancement via Generalized Eigendecomposition*** (GED)^35,52^. We first applied GED to hippocampal source time series to identify spatial filters that maximise theta-band variance relative to broadband activity. GED enhances certain features of the data defined by the experimenter, by creating spatial filters that maximise specified contrasts. For our case, it was important to enhance the signal in the theta frequency band. To achieve that, a narrowband covariance ***S*** was computed from theta-band filtered data and a broadband reference covariance ***R*** was computed from the unfiltered data:

- *R = cov(X)*, where *X* is the hippocampal source matrix, samples x sources (no band-pass)
- *S = cov(X_0_),* where *X_0_* is *X* band-pass filtered in 2-8 Hz, sing a zero-phase finite impulse response (FIR) filter, implemented via FFT-based convolution with symmetric edge trimming. We then solved the generalized eigenproblem:

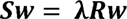

where eigenvectors *w* define spatial filters and eigenvalues λ quantify the ratio of theta-band to broadband power. The leading eigenvector (w_1_, associated with the largest λ) was retained as the spatial filter that maximises theta specificity. The corresponding component time course was reconstructed by linearly projecting the broadband unfiltered data through w*_1_*. This procedure reduced hippocampal sources to a single component optimally tuned to theta-band activity. Crucially, GED does not band-pass the data itself: the filter is derived from band-limited covariance matrices but applied to the broadband signal. The resulting component is a weighted sum of hippocampal sources that preserves native waveform features while enhancing theta contributions.
3. ***Theta frequency identification using the cyclic homogeneous oscillation method*** (CHO)^36^. To identify transient oscillatory bursts, we applied the CHO method to the GED-derived hippocampal component time series. CHO tests candidate oscillations against three criteria: (i) events must exceed the 1/f aperiodic background in both time and frequency domains; (ii) events spanning fewer than two complete cycles are excluded; and (iii) autocorrelation is used to verify periodicity, distinguishing genuine oscillations from harmonics or spurious events. An advantage of CHO is that oscillations are verified via autocorrelation, so the algorithm does not assume sinusoidal shape (which is not always the case for genuine oscillations). For each event passing these criteria, CHO returned onset and offset times, frequency range, centre frequency, and number of cycles. Only events with centre frequencies in the theta range (3–7 Hz) were retained.

The same pipeline parameters were applied across all datasets unless otherwise noted.

### Rodent dataset

We analysed 64 intracranial electrophysiological recordings from the entorhinal cortex of rodents (*N* = 3) during open-field navigation (data from Ref ^53^). Local field potential (LFP) and locomotor speed signals were processed in MATLAB (R2023b, MathWorks, Natick, MA, USA). Continuous data were segmented into epochs of 1 second, and the average speed was computed for each segment.

To obtain the theta covariance matrix required for GED, LFP signals were finite-impulse-response (FIR) filtered between 2–15 Hz. In 9 of the 64 sessions, only a single LFP channel was available. In these cases, the GED step was not performed, and the CHO method was applied directly. For subsequent analyses, only frequencies between 6 and 10 Hz were retained, consistent with other studies investigating the same relationship.

To test the relationship between theta frequency and running speed, we computed Pearson correlation. To limit the influence of tracking artefacts, epochs with locomotor speeds > 5 SD from the session mean were excluded (<0.1% of epochs). Importantly, results were unchanged whether or not these outliers were retained. To account for repeated measures across animals and sessions, we used linear mixed-effects models (lme4 package in R). Locomotor speed was entered as the dependent variable, theta frequency as the fixed effect, and random intercepts were specified for rodent identity and recording session, using Wilkinson notation:

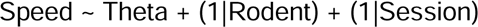

### 4-Hz RSS MEG dataset

Full details about the material and methods relative to this dataset are reported in Wang et al. (2025)^54^. Here, we report only the relevant details for the current analysis. Twenty-four healthy participants were included in this analysis (mean age: 24.1 years; range: 18–35 years; 14 females).

#### Task description

The task consisted of an associative memory paradigm with an encoding, a distractor, and a retrieval phase (Fig. 2b).

During encoding, participants were presented with pairs of unrelated videos and sounds. Stimulus pairs were displayed for 3 seconds and were modulated at 4 Hz. Stimuli were presented either in synchrony (0° phase offset; synchronous condition) or out of phase (asynchronous condition, where the sound was shifted by 180° from the video). To account for differences in sensory processing latencies, the auditory stimulus presentation was always delayed by 40 ms relative to visual onset. On each trial, participants were asked to rate how well the sound and the video fit together on a scale ranging from 1 = “not at all” to 5 = “very well”, to ensure active associative encoding. The encoding phase was followed by a distractor task, in which participants judged whether visually presented numbers were odd or even, preventing rehearsal. During retrieval, participants were cued with a previously encoded sound and asked to select the matching video from four still frames, providing a forced-choice measure of associative memory. Participants completed 12 blocks of 16 trials each, for a total of 192 trials evenly divided between conditions (96 synchronous, 96 asynchronous).

#### MEG data acquisition and preprocessing

Full details of online data acquisition are reported in Wang et al. (2025)^54^. In brief, brain activity was recorded using a whole-head 306-channel MEG system (TRIUX Neo, MEGIN Oy, Espoo, Finland), with 204 planar gradiometers and 102 magnetometers. All preprocessing steps were performed using MNE-Python^55^. Signals were band-pass filtered between 1 and 148 Hz, and visually identified bad channels were excluded. Environmental noise was attenuated using temporal Signal-Space Separation (tSSS) with Maxwell filtering, incorporating continuous head-position tracking to correct for head movements. Line noise was removed with notch filters at 50 and 100 Hz. Muscle artefacts were detected by z-scoring the signal filtered between 110 and 140 Hz, and segments exceeding a threshold of 10 were discarded. Data were downsampled to 500 Hz for independent component analysis (ICA). Eyeblink- and heartbeat-related components were identified based on their topographies and time courses and removed from the original data (non-downsampled). Finally, data were segmented into epochs of 7 seconds (–2 to +5 s relative to the stimulus onset) and visually inspected. Trials containing residual artefacts were excluded.

#### Source reconstruction

Source reconstruction was performed in MNE-Python. We constructed the forward model using a single-shell boundary element model (BEM), derived from each participant’s anatomical 3T MRI segmented with FreeSurfer. The source space included a 5-mm volumetric grid covering the bilateral hippocampi and a coarsely sampled cortical surface. This approach allowed higher spatial resolution in the hippocampi while maintaining computational efficiency for whole-cortex coverage. We performed MEG–MRI coregistration using digitised fiducial landmarks and scalp points, and from these elements computed the lead field matrix describing the mapping between dipole sources and MEG sensors.

To reconstruct source activity, we applied a linearly constrained minimum variance (LCMV) beamformer spatial filter. The data covariance matrix was whitened using empty-room recordings (5 minutes collected before each experimental session), which underwent the same preprocessing as the experimental data, to avoid amplification of noise components when combining magnetometers and gradiometers. To account for the rank deficiency of the data, 5% of the sensor power was used to apply a regularisation (Tikhonov) to the covariance matrix. Source orientations were optimised to maximise output power, and unit-noise-gain normalisation was applied to mitigate the bias toward superficial sources ^51,56^. The resulting spatial filters were applied to the experimental data to reconstruct time series for each grid point.

#### Frequency estimation

The frequency analysis was performed in MATLAB, as GED and CHO are already implemented and validated in this environment. Trials were segmented into 1-s chunks, resulting in 7 time bins per trial (–2 to 0 s pre-stimulus, 0 to 3 s stimulation, 3 to 5 s post-stimulus). For each chunk, GED was applied to enhance theta-band content. The covariance matrix of interest was computed from data filtered to 2–8 Hz and contrasted with the broadband covariance matrix; the first GED component was retained and projected back onto the data to obtain the time series. The CHO method was then applied to detect oscillatory events and estimate their centre frequency. Only frequency values between 3 and 7 Hz were stored.

#### Statistics

Statistical analyses were performed in RStudio^57^ (v4.3. 2) on subject-level averages. For each participant, frequency deviation was averaged across trials separately for each source-phase condition and time window. These values were entered into a repeated-measures ANOVA with within-subject factors Condition (0° vs 180°) and Time (7 levels). Sphericity was assessed using Mauchly’s test and was not violated. Effect sizes are reported as generalized eta squared (ges). Post-hoc comparisons across time bins were performed using estimated marginal means, and p-values were corrected for multiple comparisons using the false discovery rate^58^ (FDR).

### MEG–iEEG dataset

We analysed 6 simultaneous MEG–iEEG recordings obtained from 2 patients with pharmaco-resistant epilepsy undergoing presurgical monitoring at Ruijin Hospital, Shanghai. One patient contributed four recordings, with depth electrodes implanted bilaterally in the hippocampus (two runs from the left hippocampus and two from the right hippocampus). The second patient contributed two recordings from the left hippocampus. Depth electrodes (Beijing Sinovation Medical Technology, Beijing, China) had 8 or 16 contacts (2-mm length, 0.8-mm diameter, 1.5-mm inter-contact spacing). Implantations were performed under general anaesthesia using the orthogonal method with a Leksell head frame. MEG data were acquired simultaneously using a whole-head Elekta Neuromag VectorView 306-channel system (102 magnetometers, 204 planar gradiometers) in a magnetically shielded room, with integrated EEG hardware for iEEG recording. Data were sampled at 1000 Hz and recorded at rest with eyes closed. The MEG data were provided with temporal signal space separation (tSSS) already applied for noise suppression. Ethical approval and written informed consent were obtained by the recruiting site.

#### Preprocessing (MEG and iEEG)

MEG was processed in MNE-Python. Signals were band-pass filtered (1–148 Hz), notch-filtered at line frequencies (50/100 Hz), visually inspected to mark bad channels, and cleaned using ICA. For the second patient, a signal dropout introduced a discontinuity in the MEG–iEEG data. To ensure stable artefact decomposition, the data were split into two continuous segments before applying ICA. Each segment was denoised independently, after which they were concatenated at the epoching step. However, for analyses that rely on component stability (e.g., CCA), the two segments were treated separately. Data were segmented into 1-second non-overlapping epochs. The same filters were applied to the iEEG channels, which were then epoched identically and restricted to hippocampal contacts. Electrode contacts were localised automatically using the AAL3 atlas; only contacts labelled as hippocampal were retained for further analyses. In the first patient, eight contacts were confirmed in the right hippocampus (but two were excluded due to bad signal) and six in the left hippocampus. In the second patient, three contacts were confirmed in the left hippocampus. All analyses were restricted to these hippocampal contacts. To minimise contamination by epileptiform activity, interictal epileptiform discharges (IEDs) were detected using a semi-automatic pipeline ^59^ based on Gelinas et al. criteria^60^. Time points containing visually confirmed IEDs were annotated, and the corresponding epochs were excluded from subsequent MEG and iEEG analyses.

#### Source reconstruction

Source reconstruction followed the same general pipeline as described for the 4-Hz RSS MEG dataset. For the MEG–iEEG recordings, the source space was restricted to the hippocampus corresponding to the one included in the recordings (e.g., only left hippocampus when electrodes were confined to the left side). This ensured spatial specificity and reduced unnecessary source dimensions. The noise covariance was estimated using the ad hoc method provided in MNE-Python (function: *mne.make_ad_hoc_cov*) as empty-room recordings were not available.

#### Frequency estimation

The same pipeline was applied in parallel to hippocampal iEEG signals (restricted to contacts anatomically localised to the hippocampus) and to MEG virtual electrodes of the same hippocampus reconstructed with the LCMV beamformer. This allowed direct comparison of frequency estimates across recording modalities. We used the same pipeline parameters as in the RSS dataset but broadened the accepted frequency range, retaining oscillations with centre frequencies between 2 and 8 Hz. Statistics

Analyses were restricted to epochs in which both pipelines returned valid frequency estimates (i.e., non-NaN in MEG and iEEG), ensuring a one-to-one correspondence between modalities. For each patient and hippocampal region, trials from all runs were concatenated and Pearson correlation was computed between matched iEEG and MEG theta frequencies.

To assess the statistical significance of these correlations, we implemented non-parametric permutation tests separately for each participant and hemisphere. Within each permutation (10,000 iterations), the correspondence between MEG and iEEG frequencies was disrupted by randomly permuting MEG values within each run, thereby preserving the within-run structure. The runs were then re-pooled, and the correlation was recomputed. The observed correlation was compared to the resulting null distribution to obtain a p-value reflecting the probability of observing an equal or greater correlation under the null hypothesis of no correspondence between modalities.

To further evaluate how faithfully MEG captured the structure of the underlying hippocampal theta rhythm, we constructed a frequency-bin confusion matrix. Single-trial theta frequencies from both modalities were binned into non-overlapping frequency intervals across the analysis band (2–8 Hz). Confusion matrices were computed with rows representing iEEG bins and columns representing MEG bins. To account for unequal occupancy across bins, matrices were row-normalised. The diagonal elements quantify the extent to which MEG preserves the coarse frequency structure present in hippocampal iEEG.

As a summary statistic, we computed a diagonal-to-off-diagonal ratio, defined as the sum of the diagonal entries (correct bin assignments) divided by the sum of all off-diagonal entries (misassignments). Statistical significance of this ratio was assessed with a run-wise permutation test analogous to the correlation analysis. In each permutation, MEG frequency labels were randomly shuffled within run, preserving the temporal and session structure of the data. A null distribution of 10,000 diagonal/off-diagonal ratios was obtained, and the observed ratio was converted into a p-value reflecting the likelihood of observing equal or stronger modality agreement by chance.

#### Canonical Correlation Analysis (CCA)

MEG sensor data after ICA (306 channels) and hippocampal iEEG data were downsampled to 50 Hz and z-scored. CCA was then applied to find linear projections of MEG and iEEG that maximised their correlation across time. We regularised this with an L2 penalty^61^, which we optimised via Bayesian Adaptive Direct Search (BADS^62^) within a nested cross-validation^63^ (5 folds). To localise the solution, we predicted the CCA component (i.e., the projection of MEG data through the CCA MEG weights) from source-level activity estimated at individual grid points using subject-specific LCMV beamformer filters (10-mm grid; all orientations combined into three-dimensional predictors per grid point). This regression was again ridge-regularised, with the penalty optimised using BADS in nested cross-validation. For each grid point, we computed the held-out test-set correlation between predicted and observed signals across folds, producing a whole-brain performance map and its peak location.

#### Cross-correlation

For the cross-correlation analysis, MEG and hippocampal iEEG data were taken from the same preprocessing stage used in the CCA analysis (after ICA cleaning and IEDs removal). MEG data were projected to source space using precomputed LCMV beamformer filters (10-mm grid, single-orientation) derived in the validation pipeline. For each grid point, the broadband MEG time series was low-pass filtered at 25 Hz, projected with the corresponding spatial filter, resampled to 50 Hz, and z-scored. Cross-correlations were then computed independently for each hippocampal iEEG electrode and each MEG grid-point time course, within a restricted lag window of ±1 s (±50 samples at 50 Hz). To avoid bias, we implemented a 5 fold cross-validation procedure: in each iteration, cross-correlations were computed on four folds to determine the optimal lag and evaluated on the held-out fold to assess generalisation. Each fold served once as the test set, resulting in five cross-validated correlation estimates per subject, which were then averaged to obtain a single cross-validated lag-sensitivity estimate for each grid point and electrode.

## Data availability

The preprocessed data from the 4-Hz RSS MEG project are available at https://osf.io/skm4q/. The rodent intracranial dataset and the MEG–iEEG dataset analysed in this study were obtained from previously published studies and are subject to the data-sharing policies of their original investigators. Access to these datasets may be obtained from the respective data owners upon reasonable request.

## Code availability

Custom code used to generate the results reported in this study is available at https://github.com/NoT-CoOLab/MEG-theta-pipeline-validation. Any additional information required to reanalyse the data reported in this paper is available upon request.

## Acknowledgements

This study was supported by the Medical Research Council Doctoral Training Program in Precision Medicine (MR/W006804/1 to E.M.). D. B. is supported by a UKRI Frontier Research grant (EP/X023060/1).

## Author contributions

Conceptualization: S.H., E.M.; Investigation: E. M., D.W., C.C., S.Z., B.S., D.B.; Data Curation: D.B., C.C., S.Z., B.S., E.M., D.W., S.H.; Software: E.M., C.D., D.B., S.P.; Formal Analysis: E.M. and C.D.; Writing – original draft: E.M., C.D., S.H.; Writing – review & editing: E.M., C.D., D.W., C.C., B.S., S.Z., R.A.A.I, L.P., S.P., D.B., and S.H.; Funding acquisition and supervision: S.H.

