## supplementary material for "Non-invasive tracking of hippocampal theta oscillations"

### Supplementary information

#### Supplementary Methods

##### 4 Hz RSS MEG study - Control ROI analysis

The same source reconstruction pipeline described in the main Methods was applied for this control analysis. The only modification was the definition of the source space: instead of restricting the beamforming grid to the hippocampus, additional volumetric masks corresponding to the primary motor cortex (M1) and dorsolateral prefrontal cortex (DLPFC) were included. These regions were defined using the standard Desikan-Killiany atlas labels available in MNE-Python (labels *ctx-rh-rostralmiddlefrontal* and *ctx-lh-rostralmiddlefrontal* for DLPFC; labels *ctx-rh-precentral* and *ctx-lh-precentral* for M1). All the other parameters regarding inverse modelling were identical to those used in the main hippocampal analysis.

For each ROI, the theta frequency was estimated every second across the full 7s trial. These per-second estimates were then averaged into windows as follows: Pre = mean of the two 1s bins preceding stimulation, Stim = mean of the three 1s bins during stimulation, and Post = the first 1s bin after stimulation, selected to capture the entrainment echo (leaving out the last second of the trial). Frequencies were expressed as deviations from the 4 Hz stimulation frequency and then baseline-corrected by subtracting the mean frequency in the Pre window. Values were averaged across the two phase-offset conditions (0° and 180°) for each participant.

#### Supplementary Results

##### 4 Hz RSS MEG study - Control ROI analysis

A one-way repeated-measures ANOVA with ROI (HIPP, M1, DLPFC) on baseline-corrected Δ values revealed a significant effect during stimulation (*Stim - Pre*: *F*(2,46) = 45.65, *p* < .001, ges= .40). Post-hoc paired t-tests (FDR-corrected) showed that frequency shifts in the hippocampus were larger than in M1 (*t*(23) = -9.19, *p* < .001) and DLPFC (*t*(23) = -7.54, *p* < .001), with no difference between neocortical ROIs (p > .69). In the post-stimulation period (Post – Pre), the ROI effect was not significant (F(2,46) = 0.06, p = .94), indicating that the regional difference observed during stimulation did not persist into the post-stimulus period.

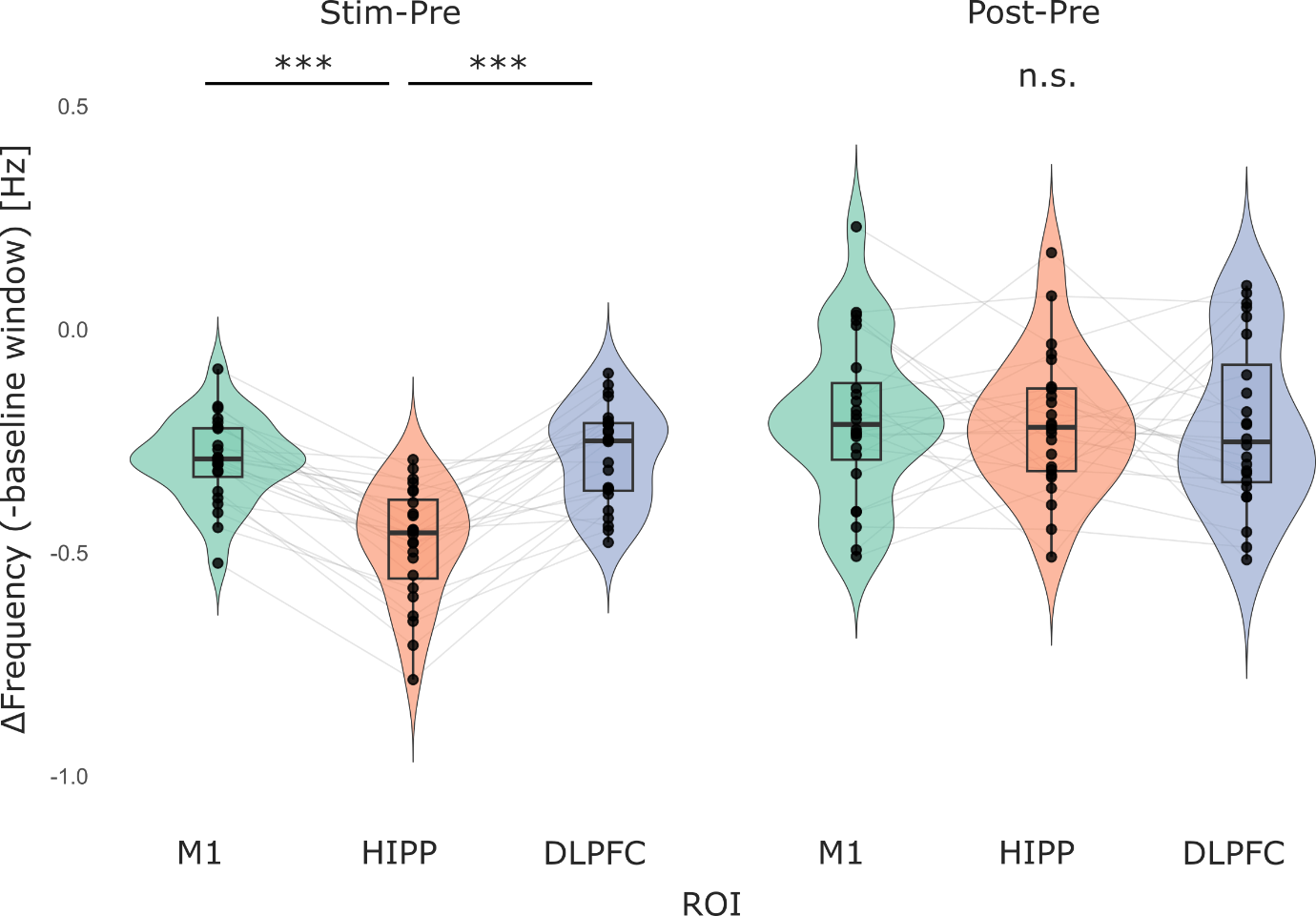

**Supplementary Fig. 1. Control analysis of baseline-corrected frequency changes across ROIs and time windows.**

Raincloud plots show individual (black dots, grey lines) and group-level (violin) changes in the identified theta centre frequency relative to the 4 Hz stimulation frequency, baseline-corrected to the pre-stimulation window, across three cortical ROIs: primary motor cortex (M1), hippocampus (HIPP), and dorsolateral prefrontal cortex (DLPFC). Two temporal contrasts are shown: the stimulation window (Stim - Pre, *left*) and the early post-stimulation window (Post - Pre, *right*).

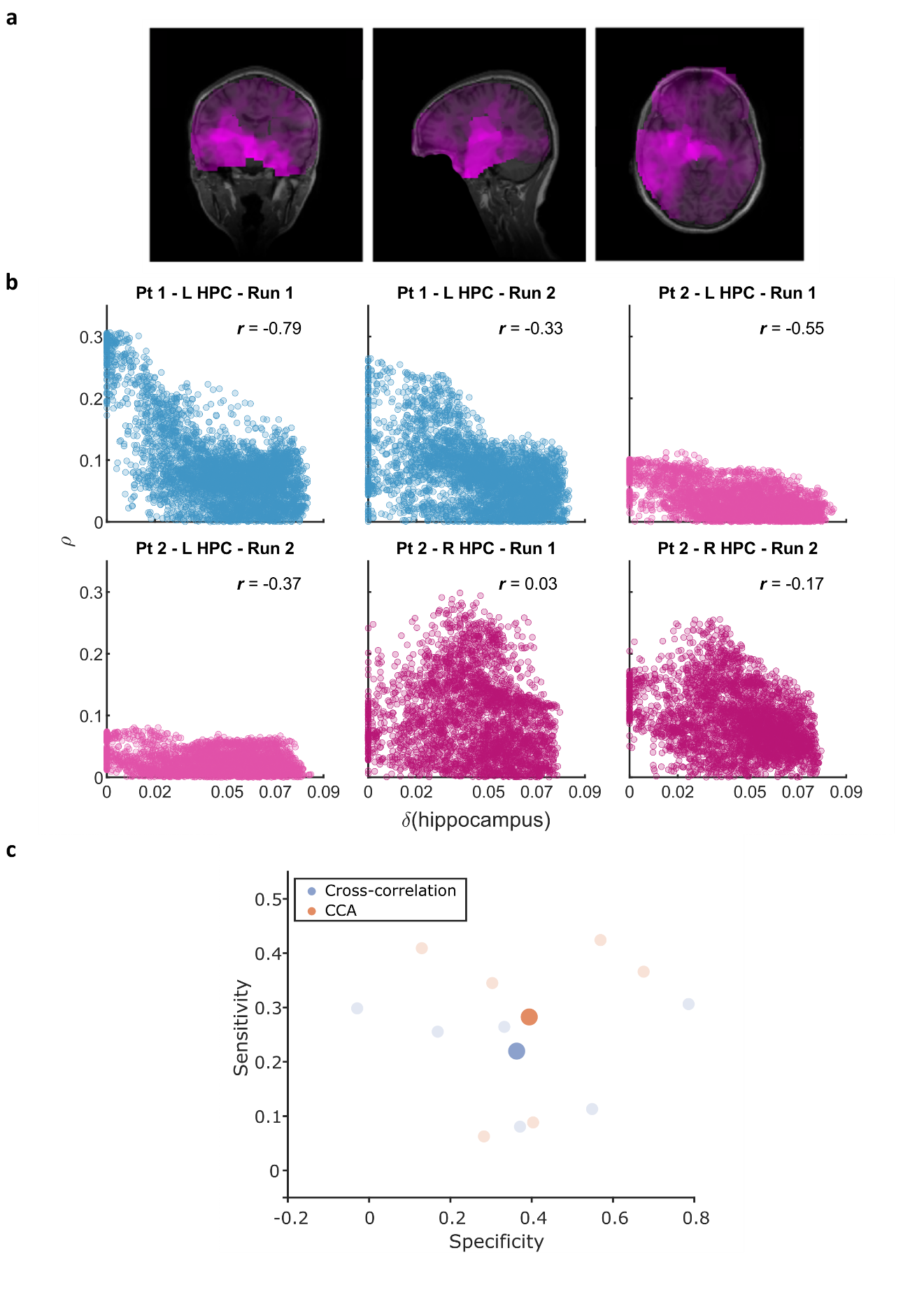

**Supplementary Fig. 2. Cross-correlation analysis on the MEG-iEEG dataset and comparison with CCA**

**a** Example of one participant (Patient 2 - left hippocampus; best performing run) illustrating the cross-correlation values for each grid point of the source space (cross-validated correlation at each grid point; blue overlay). **b** Cross-correlation performance (best electrode) as a function of hippocampal distance per participant and run. Each scatter shows one run. Each point is a source-space grid location. The x axis shows Euclidean distance to the nearest hippocampal voxel (δ), the y axis shows the cross-validated model performance (p). Point colour indicates the lag (s) at which the peak correlation occurred. Numbers report the Pearson correlation between δ and p, used as a specificity index (negative values denote stronger hippocampal focus). **c** Sensitivity versus specificity across participants and runs. CCA shows higher overall sensitivity than cross-correlation while maintaining comparable specificity. Each point represents one run; large circles indicate group means.

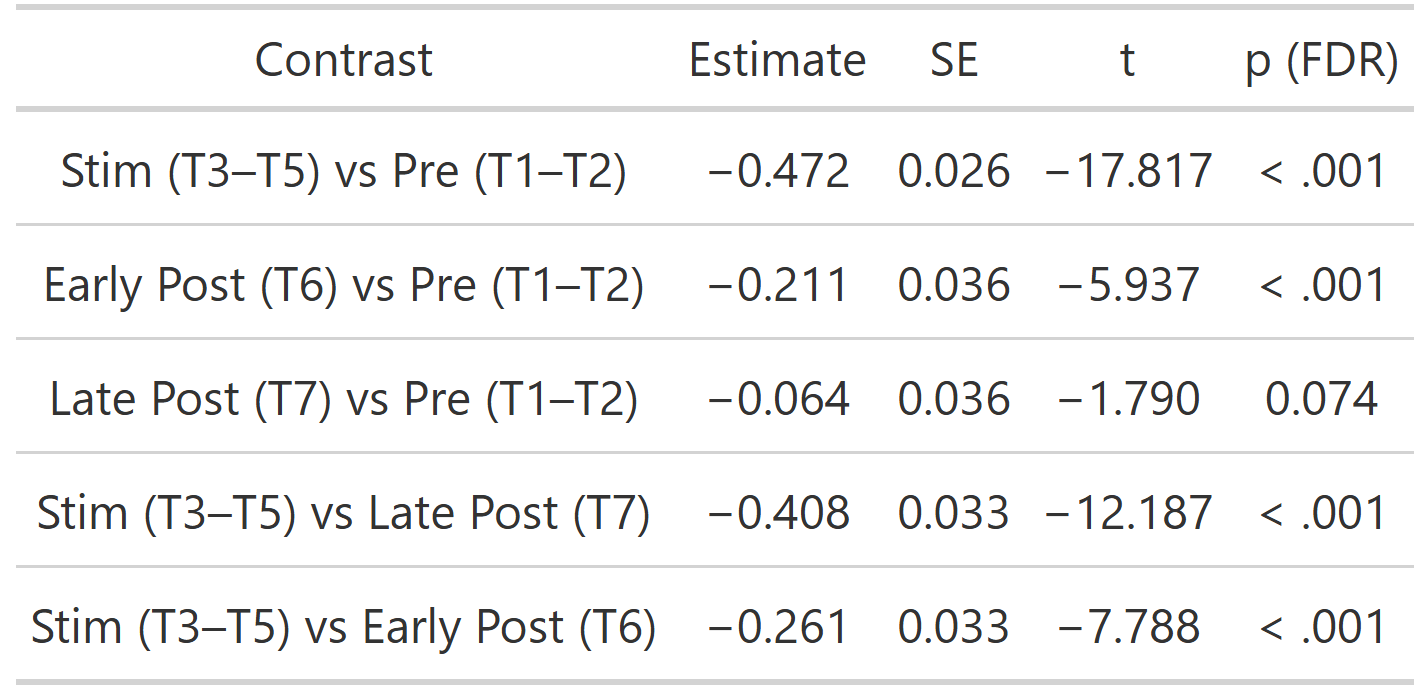

**Supplementary Table 1. Planned contrasts derived from estimated marginal means.**

Pre corresponds to time bins 1–2, Stim to bins 3–5, Early Post to bin 6, and Late Post to bin 7. P-values were corrected for multiple comparisons using the false discovery rate (FDR).
